# Artificial immune cell, *AI-cell*, a new tool to predict interferon production by peripheral blood monocytes in response to nucleic acid nanoparticles

**DOI:** 10.1101/2022.07.28.501902

**Authors:** Morgan Chandler, Sankalp Jain, Justin Halman, Enping Hong, Marina A. Dobrovolskaia, Alexey V. Zakharov, Kirill A. Afonin

**Author notes:** these authors contributed equally to the project (Morgan Chandler, Sankalp Jain). To whom correspondence should be addressed: Kirill A. Afonin,; Alexey Zakharov,; Marina A. Dobrovolskaia.

## Abstract

Nucleic acid nanoparticles, or NANPs, are rationally designed to communicate with the human immune system and can offer innovative therapeutic strategies to overcome the limitations of traditional nucleic acid therapies. Each set of NANPs is unique in their architectural parameters and physicochemical properties, which together with the type of delivery vehicles determine the kind and the magnitude of their immune response. Currently, there are no predictive tools that would reliably guide NANPs’ design to the desired immunological outcome, a step crucial for the success of personalized therapies. Through a systematic approach investigating physicochemical and immunological profiles of a comprehensive panel of various NANPs, our research team has developed a computational model based on the transformer architecture able to predict the immune activities of NANPs *via* construction of so-called artificial immune cell, or *AI-cell*. The *AI-cell* will aid addressing in timely manner the current critical public health challenges related to overdose and safety criteria of nucleic acid therapies and promote the development of novel biomedical tools.

## INTRODUCTION

Therapeutic nucleic acids (TNAs) have enriched and diversified the landscape of nanomedicine^1^, and their clinical success brought about the development of novel biomolecular platform, based on nucleic acid nanoparticles, called NANPs^2, 3^. NANP technologies aim to advance the programmability of TNAs, tune their physicochemical and biological properties, and optimize the formulation and storage processes. The bottom-up assembly of NANPs takes advantage of nucleic acids’ folding pathways along with the several computational tools available for precise coordination of sequence design and expanded repertoire of structural and interacting motifs^4-7^. Hundreds of NANPs have been engineered to vary in chemical composition, sizes, and shapes that range from three-dimensional assemblies down to linear nanoscaffolds. Individual oligonucleotides in NANP compositions may be additionally defined in certain lengths and GC content, while also incorporating various TNAs (*e*.*g*., siRNA, aptamers, anti-sense oligonucleotides, CpG DNAs), proteins, small molecules, and imaging agents suitable for biomedical applications^8-12^. Consequently, a growing library of functional NANPs have been shown to operate in response to other classes of biomolecules, or stimuli while executing therapeutic decisions based on the environmental inputs^13-15^.

While the practicality of NANPs offers new ways to treat a broad spectrum of malignancies that span from cancers to infectious and cardiovascular diseases^16^, the intended clinical applications and routes of administration place NANPs’ interactions with the human immune system to be carefully considered and understood for further translation of this technology into the clinic^17, 18^. The immune recognition of these novel nanomaterials is inherent to the natural line of immune defense evolved for the detection of nucleic acids associated with pathogen invasion and cellular damage^4, 19-22^. However, their unique architectural parameters and chemical compositions define the NANPs’ immunorecognition which cannot be extrapolated from the immune responses to pathogen- or damaged self-associated nucleic acids and conventional TNAs^19^. The ability to predict how NANPs interact with the human immune system would allow for tailoring their formulations to the specific biomedical task with maximized therapeutic effects and controlled immunological activity, which collectively are required to achieve desired therapeutic efficacy and safety. In addition, as was revealed by our recent studies^8-10, 19, 23-28^, NANPs can function not merely as nanoscaffolds for TNAs but also as independent immunostimulatory therapeutics with conditional intracellular activation of intended functions beneficial for vaccines and immunotherapies. For over 10 years, our team created a comprehensive library of NANPs, designed by our group and others, and subjected them to detailed physicochemical characterization, sterility and endotoxin assessments, and immunological assays carried out in model cell lines and in primary human peripheral blood mononuclear cells (PBMCs)^24^. PBMCs were chosen as the most accurate pre-clinical model that produces the most predictive results for cytokine storm toxicity in humans^29^.

Translating NANP materials from bench to the clinic requires quick coordination of design principles. The incorporation of a particular level of immunostimulation and matching it to the desired application requires feedback from the experimental analysis to the computational design phase, which in turn entails complete recharacterization of NANPs and can delay their production. To improve this pipeline, several design parameters based around a representative set of NANPs have been previously correlated with cytokine production in model cell lines to determine trends of the immune response^27^. Moreover, deep learning contributed to major advancements in several research fields ranging from computer vision to natural-language processing. It is also widely applied in biomedical research areas such as drug discovery and genomics^30^. In genomics, sequence-based deep learning models outperformed classical machine learning^31^ and also enabled efficient prediction of the function, origin, and properties of DNA and RNA sequences by training neural networks on large datasets^32-37^. A robust model that can predict immune responses will have an enormous benefit in the design of NANPs. Our earlier quantitative structure-activity relationships (QSAR) modeling utilized a dataset collected for 16 NANPs which were assessed in model cell lines, and demonstrated that computational prediction of experimentally observed immunomodulatory properties is feasible^27^.

Despite this progress, there is currently no reliable bioinformatics tool to computationally identify optimal NANPs structure and match it to the desired immunological outcome. Such tool would tremendously accelerate NANPs design and selection for personalized immunotherapeutic approaches or immunologically safe nanoscaffolds for other indications in which the stimulation of the innate immune responses is not wanted. Therefore, our present study was conducted to improve the communication between biotechnology, immunology, and bioinformatics and to create a new tool which would enable a prediction of NANPs structure-activity relationships in order to better guide the overall designing principles (**Figure 1A**). Particularly, we employed random forest (RF) and two different neural network architectures (a recurrent neural network, a transformer neural network) to develop models that predict immunomodulatory activity for a much larger set of 58 NANPs that have been uniformly characterized using previously established, clinically relevant models^24^. Long-short-term memory (LSTM) architecture was used as the recurrent neural network. While the RF models use physicochemical properties derived from the constructed nanoparticles, the neural networks learn directly from the nanoparticle sequences. The neural network architectures investigated in this study facilitate discovery of hidden patterns *via* non-linear transformation of raw sequence data. These methods may also be applied to designing new NANPs (**Figure 1B**).

**Figure 1.**
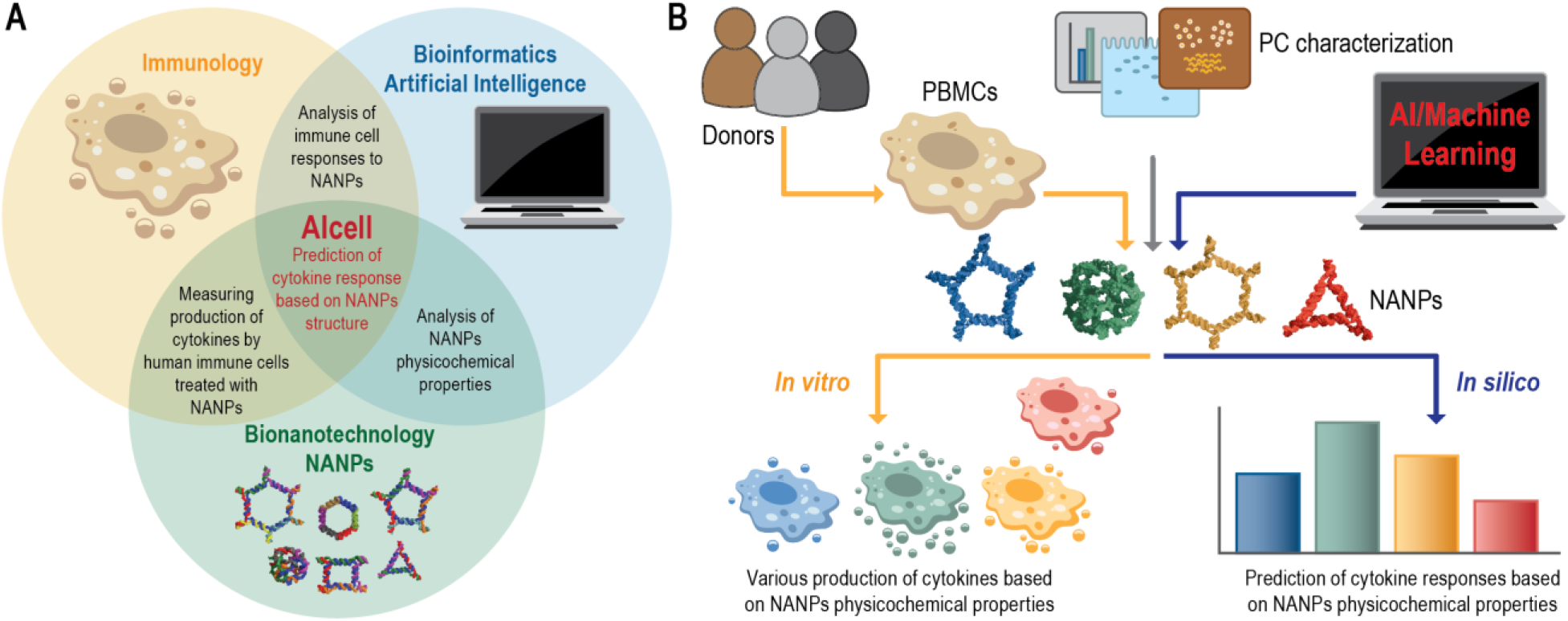
Conceptual representation of artificial immune cell (*AI-cell*) tool. (**A**) The initial design and synthesis of nucleic acid nanoparticles (NANPs) is followed by their physicochemical characterization and assessment of immunostimulatory potential to then be applied for predictive computational analysis of the NANPs immune responses. (**B**) The experimental workflow used for the *AI-cell* development.

The top performing models resulted from this study are freely accessible to the research community *via* (**https://aicell.ncats.io/**) and can be applied to predict the immunological responses of any novel nucleic acid architecture.

## RESULTS

### Representative NANP Database

We designed a library of representative NANPs to study key structure-activity relationships that define NANP interactions with the cells of the human immune system (**Figure 2** and **SI Table 1**). Our dataset includes 58 different functional and non-functional NANPs made of either DNA or RNA, having planar, globular, or fibrous structures, different sizes, flexibilities, thermodynamic stabilities, and connectivity rules. Some of these datasets have already been published^8, 9, 19, 28, 38, 39^ (also itemized in the **SI Table 1**), whereas others were newly generated to support the development of the current AI algorithm.

**Figure 2.**
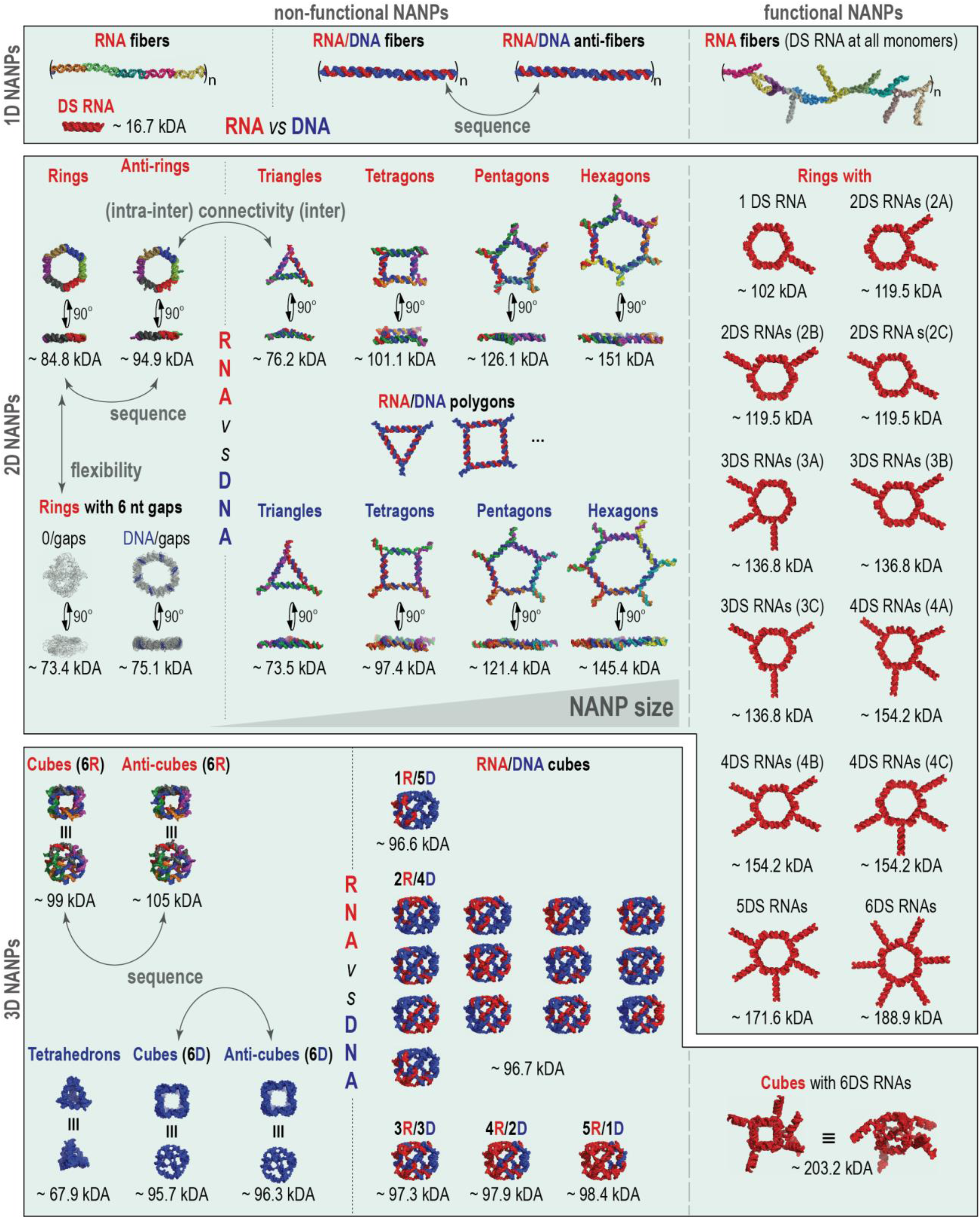
Library of representative NANPs chosen to collectively address the influence of their physicochemical properties and architectural parameters on their immunorecognition in human PBMC to further the ISAIC development.

To study the influence of architectural parameters, the immune responses to 1D fibers were compared to 2D planar structures and to 3D globular NANPs, designed by two different approaches that define the connectivity of NANPs. The first approach, represented by RNA/DNA fibers and all polygons and cubes, relies solely on intermolecular canonical Watson-Crick interactions with all NANP sequences designed to avoid any intramolecular structures^40, 41^. These design principles are characteristic for DNA nanotechnology and DNA origami^42, 43^ and allow for any RNA strand in NANP’s composition be substituted with DNA analog. The second approach, called tectoRNA^44, 45^, is exemplified by RNA rings and fibers that employ naturally occurring structural and long-range interacting motifs (*e*.*g*., kissing loops) that are rationally combined, similarly to Lego® bricks, to achieve a topological control in bottom-up assembly of NANPs ^41^.

To study the role of chemical composition, origami-like RNA NANPs were compared to their DNA and RNA/DNA analogs. This compositional blend allowed for changes in NANPs’ physicochemical and biological properties in a highly predictable and controlled manner. For example, the responses of individual NANPs to heating become different (T_m_∼36°C of the DNA cube vs. T_m_∼55.5°C of the RNA cube ^19^) and a new version of Hyperfold ^46^ can accurately predict the experimental results^8^. The chemical makeup would also influence the relative chemical stabilities of NANPs in blood serum and towards degradation by different nucleases^8, 19, 23, 38^.

To assess the effect of structural flexibility, we included gapped ring structures^47^ and cubes with different number of single-stranded uracils at their corners^19, 48^, all designed to control the dynamic behavior of 2D and 3D NANPs, respectively. Both experimental results and MD simulations supported the notion that the stability and dynamicity of NANPs can be modulated by changing the number of single-stranded regions in their structures^28, 48^.

The effect of functionalization was assessed *via* the addition of Dicer substrate (DS) RNAs^49^ to the 1D, 2D, and 3D NANPs and for the size contributions, different polygons^27^ were compared. The sequence effects were studied through the inclusion of several reverse complement structures (denoted as “anti-“) for 1D^50^, 2D^8^, and 3D^8^ NANPs. All physicochemical properties of NANPs have been characterized under the equivalent conditions and their relative immunostimulation was assessed in PBMC isolated from fresh blood drawn from healthy donors with at least three donors per each NANP. All data have been combined in a single dataset shown in **SI Table 1** with all sources for experimental results cited.

### IFN Modeling Results

With a diverse library of NANPs in the dataset, three different methods were employed to build models that predict the immunological activities of NANPs in PBMCs (**Figure 3**). First, a Random Forest (RF) method was applied using the physicochemical descriptors derived from the input sequences. The physicochemical properties of the studied NANPs along with their immune responses are provided in **SI Table S1**. Next, the neural network architectures LSTM and transformers were applied that directly learn on the NANP sequences. In both neural network models, the first step involves tokenization of the input sequences. Tokenization was performed using the ‘K-mer’ representation (K=3), which is usually employed for nucleotide sequences and the encoded sequences can be loosely considered as codons. Generating all possible combinations of 3-mers of the individual nucleotide bases (A, T, G, C, U) resulted in a total of 125 codon combinations or tokens. Each model was evaluated using five-fold cross-validation (5-CV) that was repeated for 10 times. **Figure 4A** and **4B** provides a comparison of the average performance (R^2^, RMSE) for different models generated in this study. According to the 5-CV results presented in **Figure 4**, the models developed using the transformer architecture significantly outperformed other models. The detailed model performance statistics can be found in **SI Table S2**.

**Figure 3.**
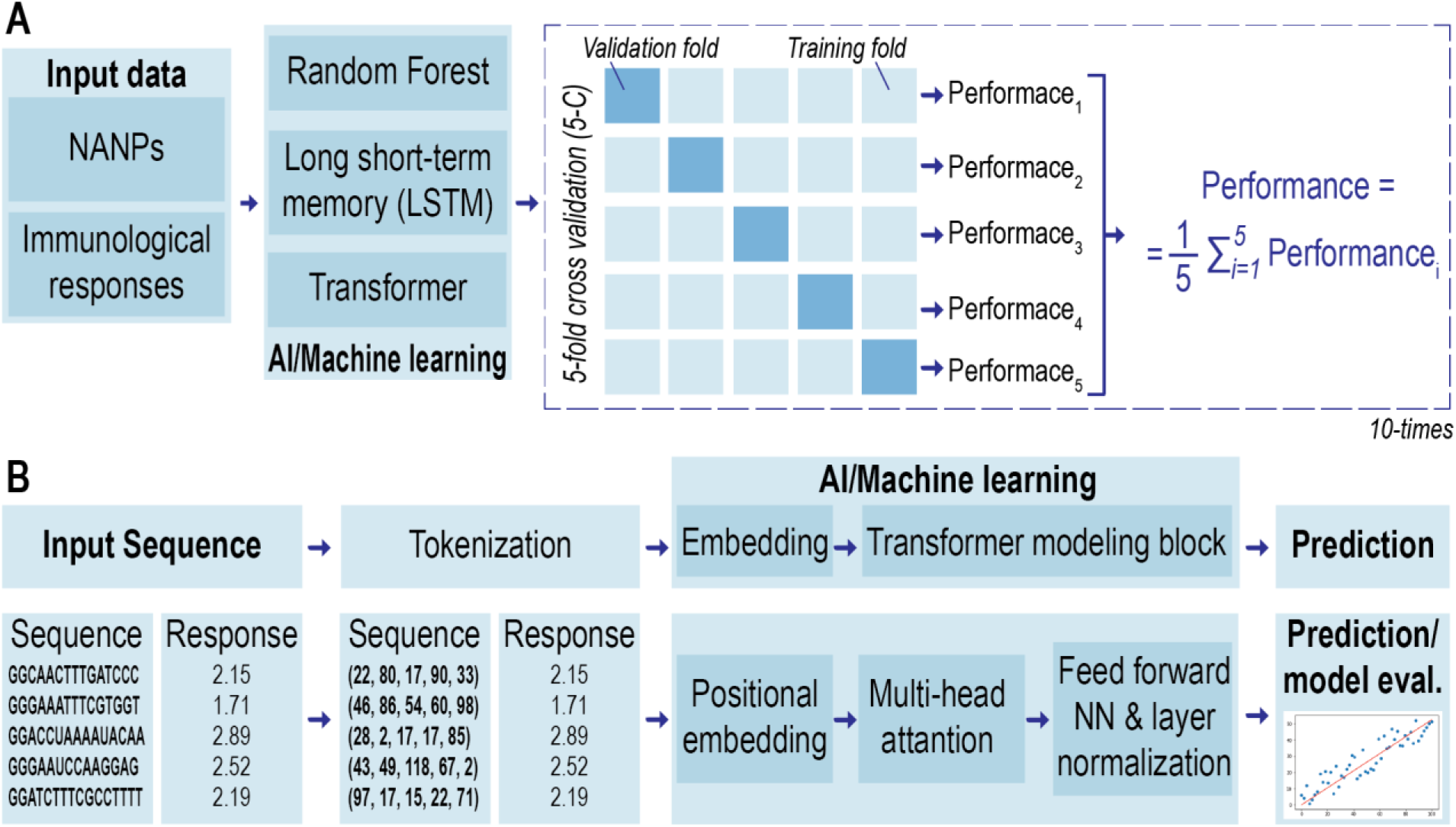
Schematic representation of the quantitative structure–activity relationship (QSAR) methodology used in this project. (A) Modeling workflow: three machine learning approaches are evaluated using five-fold cross-validation (5-CV) repeated for 10 times. (B) Overall workflow and the training procedure for prediction of nanoparticle sequence using transformer-based approach: tokenization, embedding followed by transformer modeling and prediction.

**Figure 4.**
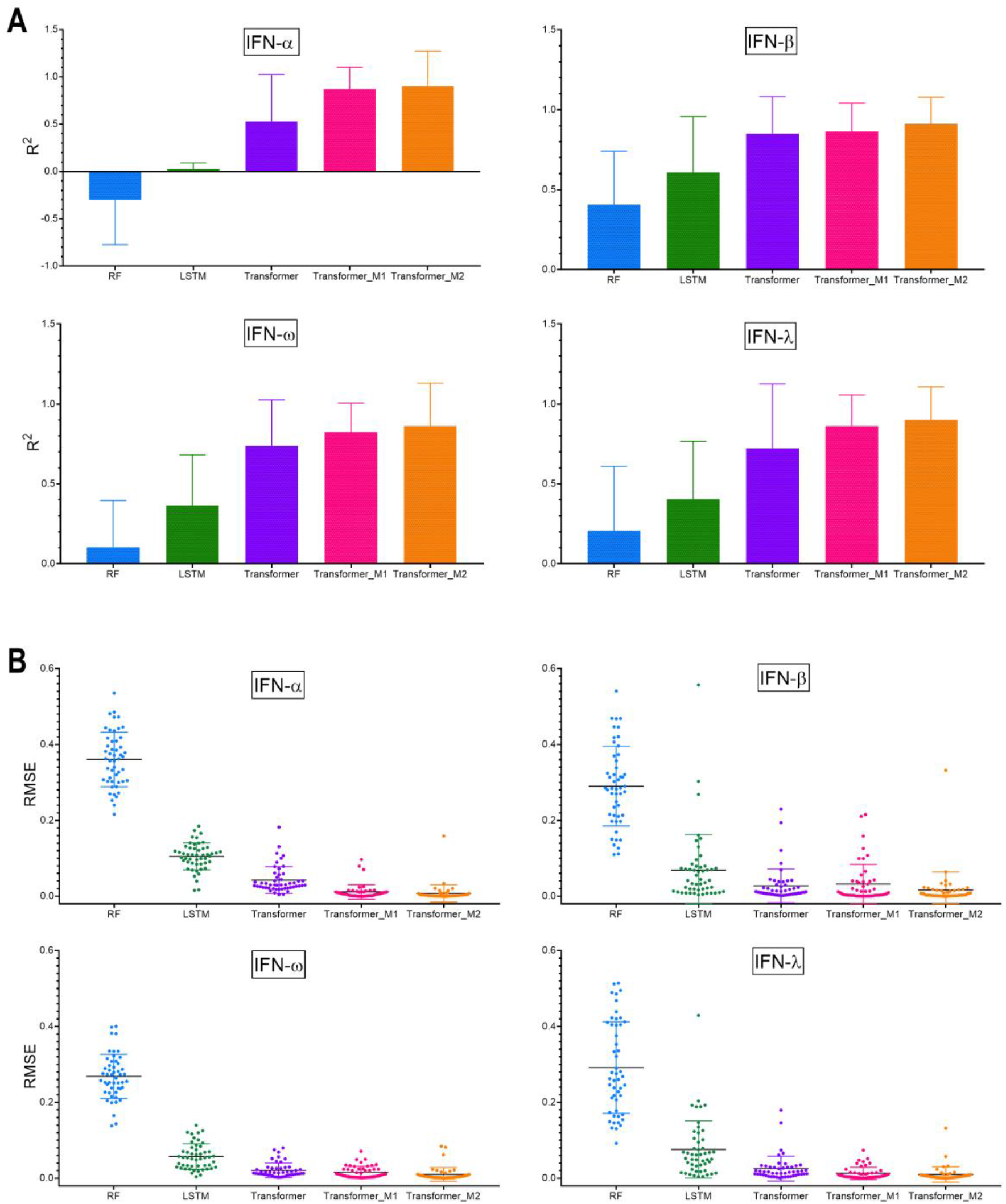
Average performance (A) *R*^2^ and (B) RMSE for different modeling approaches over 5-fold cross-validation and repeated 10-times. The error bar represents the standard deviation of the average performance over 5-folds cross-validation repeated 10-times.

Each NANP is represented by multiple sequences or strands: for example, dA, dB, dC, dD, dE, and dF are the individual sequences that form DNA cube. By default, these sequences were joined in a sequential manner for the purpose of learning. However, it is uncertain if this is the only possible configuration (*i*.*e*., arrangement of sequences) for DNA cube. To avoid any bias during learning, the best performing models based on the transformer architecture were evaluated on all possible combinations based on the different strands present in each nanoparticle. This data augmentation step led to a significant increase in the training dataset size. The models generated using this approach are referred to as Transformer_M1. The model performance improved, particularly in the case of IFN-α, where the R^2^ improved from 0.53 to 0.87. Using this approach (Transformer_M1), all models provided an R^2^ > 0.80 and RMSE < 0.03 (**Figure 4**).

It is also known that the physicochemical properties of NANPs are important for their immune response^27^. Therefore, to evaluate the contribution of the physicochemical properties to model performance, we pursued a third approach in which the physicochemical properties were combined with the sequence data to build sequence-based models. To achieve this, the numerical descriptors were converted into categories and added as tokens to the vocabulary. Tokenization was performed in the same manner on triplets. The models generated using this approach are referred to as Transformer_M2. As shown in **Figure 4A-B**, inclusion of the descriptors to the sequence-based models further improved the model performance (R^2^ > 0.85). The best prediction performance was obtained using a model that combined physicochemical properties together with sequence-based models. As seen from **Figure 4A-B**, the sequence information alone demonstrated high predictivity using transformer models (Transformer_M1), especially for IFN-β responses, and thus might play a significant role in predicting the behavior of polygons with more diverse shapes and structures.

## DISCUSSION

Machine learning and artificial intelligence have been increasingly applied in various domain such as computer vision^51, 52^, natural language processing^53-55^, drug discovery^56, 57^, QSAR^58-60^, and genomics^61-63^. AI methods such as convolutional neural networks (CNNs)^64^ and recurrent neural networks (RNNs)^65^ that are extensively used in computer vision and natural language processing have been investigated for identifying protein binding sites in DNA and RNA sequences, and achieved state-of-the-art performance^66-68^. More recently, transformer neural networks were reported to provide superior performance in the field of drug discovery and QSAR^69-71^ and demonstrated state-of-the-art results on neural machine translation task^72, 73^ including direct and single-step retrosynthesis of chemical compounds^74^. The Transformer architecture incorporates the mechanism of self-attention together with positional embedding^75^, which makes them heavily successful in the field of NLP tasks^76, 77^. Transformer-based models have also been effective in predicting novel drug–target interactions from sequence data and significantly outperformed existing methods like DeepConvDTI^78^, DeepDTA^79^, and DeepDTI^80^ on their test data set for drug–target interaction (DTI) ^81^. Another attractive task that remarkably benefits from the transformer architecture is generative molecular design. It was recently shown that transformer-based generative models demonstrated state-of-the-art performance when compared to previous approaches based on recurrent neural networks^82^. Additionally, a recent study demonstrated the application of a transformer architecture is development of a SMILES canonicalization approach that extracts information-rich embeddings and exposes them for further use in QSAR studies^83^; however the applicability of this approach to therapeutic nucleic acids and NANPs is unknown. Given the importance of nanoparticles in the field of drug delivery and the ability of NANPs to act as active pharmaceutical ingredients; offering innovative therapeutic strategies and overcoming the limitations of traditional nucleic acid therapies and the lack of predictive tools that would reliably guide NANPs design to the desired immunological outcome, we adopted transformer-based models to predict the immunological activities of the nanoparticles. An earlier study by Johnson et al.^27^ reports the use of random forest (RF) based QSAR models; however, considering that the nature of input data is sequence/text, transformer neural networks are able better learn the patterns within the data in comparison to other methods used in this study such as random forests. To the best of our knowledge this is the first study to evaluate and implement the use of state-of-the-art transformer neural networks to predict immunological activity and thus advance the current understanding of the nanoparticle properties that contribute to the observed immunomodulatory activity and establish corresponding designing principles. Our results demonstrate the benefit (significant improvement in prediction statistics; R^2^ and RMSE) of using transformer framework that is solely based on sequence data vs. RF models (**Figure 4, SI Table S2**). The data augmentation (Transformer_M1) led to further increase of the model performance. In case of QSAR modeling, the importance of data augmentation has been shown critical for the Transformer models to achieve its high performance^83, 84^. Transformer models extract semantic information in natural language processing (NLP) tasks by jointly conditioning on both left and right contexts in all layers^73^. This is particularly an essential feature in context to biological sequences, which are multidirectional in nature. The inclusion of robust sequence embeddings facilitated the proposed models to score well with the performance metrics (**Figure 4**). We expect this hybrid architecture will be continually explored for the purpose of studying NANPs.

In summary, we applied a systematic approach to connect physicochemical and immunological properties of comprehensive panel of various NANPs and developed computational model based on the transformer architecture. The resulting artificial immune cell, or *AI-cell*, predicts the immune responses of NANPs based on the input of their physicochemical properties. This model overcomes the limitations of previous QSAR model and is imperative for executing a timely response to critical public health challenges related to drug overdose and safety of nucleic acid therapies by streamlining the selection of optimal NANPs design for personalized therapies.

## METHODS

### Preparation of NANP training set

All sequences of tested NANPs are available in **SI Table S1**. A database was compiled from previously published NANPs adhering to standard methods of characterization as described below. For each NANP, the sequences of all strands included in the assembly along with the composition (DNA or RNA), quantity, and length (nts) of each strand were recorded. For each fully assembled NANP, the overall composition (DNA, RNA, or hybrid of the two), mass (g/mol), GC content (%), total number of strands in the assembly, number of helices in the structure, number of single-stranded bases, number of RNA bases, number of DNA bases, dimensionality (1D, 2D, or 3D), connectivity (origami or tectoRNA^45^), diameter (nm), melting temperature (° C), and production of IFN-α, IFN-β, IFN-ω, and IFN-λ (pg/mL) were denoted.

### NANP preparation

All DNA sequences were purchased from Integrated DNA Technologies, Inc. All RNA sequences were purchased as DNA templates and primers which were PCR-amplified via MyTaq™ Mix, 2x (Bioline) and purified using DNA Clean & Concentrator® (Zymo Research) for the preparation of double-stranded DNA templates containing a T7 RNA polymerase promotor. Templates underwent in vitro transcription with T7 RNA polymerase in 80 mM HEPES-KOH (pH 7.5), 2.5 mM spermidine, 50 mM DTT, 25 mM MgCl_2_, and 5 mM each rNTP for 3.5 hours at 37 °C and was stopped with the addition of RQ1 RNase-Free DNase (Promega, 3u/50 µL) for 30 minutes at 37 °C. Strands were purified via denaturing polyacrylamide gel electrophoresis (PAGE, 8%) in 8 M urea in 89 mM tris-borate, 2 mM EDTA (TBE, pH 8.2) run at 85 mA for 1.5 hours. Bands in the gel were visualized by UV shadowing, cut, and eluted overnight in 300 mM NaCl, TBE at 4 °C. Precipitation was performed in 2.5 volumes of 100% EtOH at - 20 °C for 3 hours, followed by centrifugation at 10.0 G for 30 minutes with two 90% EtOH washes between 10 minute centrifugations at 10.0 G. The pelleted samples were dried in a CentriVap micro IR vacuum concentrator (Labconco) at 55 °C. Pellets were dissolved in HyClone™ Water, Molecular Biology Grade (Cytiva) and concentrations were determined by measuring the A260 on a NanoDrop 2000 (ThermoFisher). NANPs were assembled in HyClone™ Water, Molecular Biology Grade (Cytiva), by adding strands in an equimolar ratio. Each NANP assembly followed previously published respective temperature steps ^8, 9, 27, 85^.

### Immunostimulation in human PBMCs

Human whole blood was obtained from healthy donor volunteers under Institutional Review Board-approved NCI-Frederick protocol OH9-C-N046. Each donor was assigned a random number. Vacutainers containing Li-heparin as an anticoagulant were used for blood collection. Research donor blood was processed to isolate PBMC within 2 hours after donation according to the protocol described earlier^86^. All NANPs were complexed with Lipofectamine 2000 (L2K) before addition to the cells as described earlier^24^. Culture supernatants were collected 24 hours after addition of NANPs-L2K, and stored at -80 °C before analysis for the presence of type I and type II I interferons using multiplex ELISA. Procedure for the interferon detection along with materials’ sources has been described earlier^24^. Some NANPs have been characterized and reported before, whereas others were synthesized and tested *de novo* to support the computational modeling of the present study (*e*.*g*., **SI Figures S1-4**). More details about new and previously studied NANPs are available in **SI Table S1**.

### Dataset for Modeling

In this study, 56 nanoparticle sequences were used to construct computational models that predict their immune responses. Based on the levels of IFN-α, IFN-β, IFN-ω and IFN-λ, four types of immune responses were identified and were used as target variables in the development of models. The complete list of the associated IFN activities and their physicochemical properties are provided in **SI Table S1**. We employed 58 NANPs to train the models; evaluated by five-fold cross-validation procedure repeated 10 times.

### Tokenization

Tokenization is considered the first step that processes the input sequence data when building a sequence-to-sequence model. It involves transformation of text input into a sequence of tokens that generally correspond to ‘words’. The nanoparticle sequences were tokenized using the K-mer representation (K=3). The K-mer representation incorporates rich contextual information for each nucleotide base by loosely encoding triplets as codons i.e., all possible combinations of 3-mers of the individual nucleotide bases (A, T, G, C, U). This resulted in a total of 125 codon combinations or tokens, which were then used to create a vocabulary. Each input sequence in the training dataset was then tokenized and passed through an embedding layer, which maps the 3-mers to vector representations.

### Generating all possible combination of the nanoparticles

It has been widely acknowledged that different nanoparticle compositions be engineered to produce desired immune responses ^8, 19^. When translating biological activity (in this case an immune response) to sequence-based learning, it is impossible to be certain about the order in which individual strands of each nanoparticle should be connected to achieve a particular immune response. Thus, to address this limitation, we generated all possible configurations using the individual strands for each nanoparticle. For a nanoparticle with ‘*n*’ strands, a total of ‘*n!*’ different combinations can be generated. IFN activity values for each combination were assigned as observed for the respective nanoparticle. This process resulted in a significantly larger training dataset. The augmented dataset was used for the training the final models. The models generated using this approach are referred to as Transformer_M1. When partitioning the augmented dataset during the 5-fold external cross-validation, to prevent the leakage of information from the training dataset to the validation dataset, all generated combination of a particular nanoparticle were either present in the training set or the validation set during. Thus, the individual cross-validation runs had no overlap of nanoparticle between the two set. In this scenario, the model statistics were calculated based on the ‘mean prediction values’ across each particle.

### Combining Physicochemical properties with Sequence-based models

Physicochemical descriptors derived from the constructures nanoparticles were previously reported to improved model performance due to their importance and relevance to the IFN activity (i.e., immune response) of nanoparticles^27^. Therefore, the physicochemical properties were used together with sequence data in development of sequence-based neural network models. For this purpose, the numerical descriptors were transformed into categories or bins (i.e., each bin encodes a value range) and added as tokens to the vocabulary previously described in the tokenization section. **SI Table S3** provides the complete list of categories for each of the eight physicochemical descriptors. Further, when generating a numerical vector for the transformer model, the tokens related to the physicochemical properties were added to the original vector after converting the input sequence to a numerical vector. The models generated using this approach are referred to as Transformer_M2.

### Modeling approaches

Two different modeling approaches were employed for the development of prediction models. In the first approach, the physicochemical properties of constructed nanoparticles were used as descriptors for creating a regression model using Random Forest (RF). RF is an ensemble of decision trees^87^ and is widely used in both classification and regression tasks. The number of trees was arbitrarily set to 100, and due to the robustness of RF^88^, no parameter optimization was performed. In the second approach, two different neural network architectures: LSTM and transformers; were employed to build prediction models that use nanoparticle sequences as input data. LSTM (Long Short-Term Memory) networks are specialized recurrent neural networks that are designed to avoid long-term dependency problem by remembering information for an extended period of time using a gating mechanism^89^. The readers are encourage to refer to the literature for further reading on LSTM networks^90^. Transformer networks have been recently introduced in the field of natural language processing^72^ and were reported to outperform recurrent neural networks architectures such as LSTM and Gated Recurrent Unit (GRU) in several NLP benchmarks on automatic speech recognition, speech translation and text-to-speech^91^. Transformers use attention mechanism and positional embeddings and facilitate encoding of multiple relationships within a sentence and process complete sentences by learning relationships between the words. The neural network architecture and the parameters used for training each of these models is provided in **SI Table S4**.

### Evaluation of model statistics

To evaluate the predictive performance of the developed models, a 5-fold external cross-validation procedure (5-CV)^92^ was employed. In this procedure, the initial data set was randomly divided into five parts. In each fold, four parts of the data were used as training set for model building and the fifth part was used as test set for assessment of external predictive performance. To be more robust and ensure that the performance obtained is not due to chance correlations, the 5-CV procedure was repeated for a total of 10 times. The performance of each model was assessed on the basis of root mean squared error (RMSE) *(Eq. 1)*, and determination coefficient R^2^ *(Eq. 2)*,

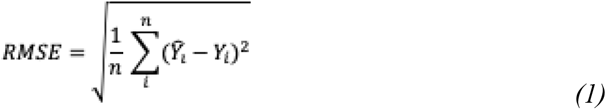

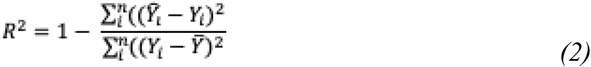

Y_i_(cap) is the predicted value for each particular sequence; Y_i_ is the observed value for each particular sequence; Y(bar) is the mean activity value from all the sequences; n is the number of sequences.

## Acknowledgements

Research reported in this publication was supported by the National Institute of General Medical Sciences of the National Institutes of Health under Award Numbers R01GM120487 and R35GM139587 (to K.A.A.). The content is solely the responsibility of the authors and does not necessarily represent the official views of the National Institutes of Health. The study was also funded in part by federal funds from the National Cancer Institute, National Institutes of Health, under contract 75N91019D00024 (M.A.D. and E.H.). This research was supported in part by the Intramural/Extramural research program of the NCATS, NIH. The content of this publication does not necessarily reflect the views or policies of the Department of Health and Human Services, nor does mention of trade names, commercial products, or organizations imply endorsement by the U.S. Government.

## Additional information

*Supplementary information*.

