## Supplementary Information for "Artificial immune cell, *AI-cell*, a new tool to predict interferon production by peripheral blood monocytes in response to nucleic acid nanoparticles"

### ABSTRACT

Nucleic acid nanoparticles, or NANPs, are rationally designed to communicate with the human immune system and can offer innovative therapeutic strategies to overcome the limitations of traditional nucleic acid therapies. Each set of NANPs is unique in their architectural parameters and physicochemical properties, which together with the type of delivery vehicles determine the kind and the magnitude of their immune response. Currently, there are no predictive tools that would reliably guide NANPs' design to the desired immunological outcome, a step crucial for the success of personalized therapies. Through a systematic approach investigating physicochemical and immunological profiles of a comprehensive panel of various NANPs, our research team has developed a computational model based on the transformer architecture able to predict the immune activities of NANPs *via* construction of so-called artificial immune cell, or *AI-cell*. The *AI-cell* will aid addressing in timely manner the current critical public health challenges related to overdose and safety criteria of nucleic acid therapies and promote the development of novel biomedical tools.

### Network architecture for LSTM models

| Layer (type) | Output Shape | Param # |
|---|---|---|
| embedding_2 (Embedding) | (None, 166, 128) | 21248 |
| lstm_3 (LSTM) | (None, 166, 128) | 131584 |
| Attention (SeqSelfAttention) | (None, 166, 128) | 16385 |
| lstm_4 (LSTM) | (None, 128) | 131584 |
| dense_3 (Dense) | (None, 200) | 25800 |
| dense_4 (Dense) | (None, 1) | 201 |

Total params: 326,802
Trainable params: 326,802
Non-trainable params: 0

### Network architecture for Transformer models

| Layer (type) | Output Shape | Param # |
|---|---|---|
| input_1 (InputLayer) | [(None, 166)] | 0 |
| token_and_position_embedding | (None, 166, 32) | 20032 |
| transformer_block (Transform | (None, 166, 32) | 10656 |
| global_average_pooling1d (Gl | (None, 32) | 0 |

```
dropout_2 (Dropout)          (None, 32)                0
_________________________________________________________________
dense_2 (Dense)              (None, 256)               8448
_________________________________________________________________
dropout_3 (Dropout)          (None, 256)               0
_________________________________________________________________
dense_3 (Dense)              (None, 1)                 257
=================================================================
Total params: 39,393
Trainable params: 39,393
Non-trainable params: 0
```

**Supporting Figures:**

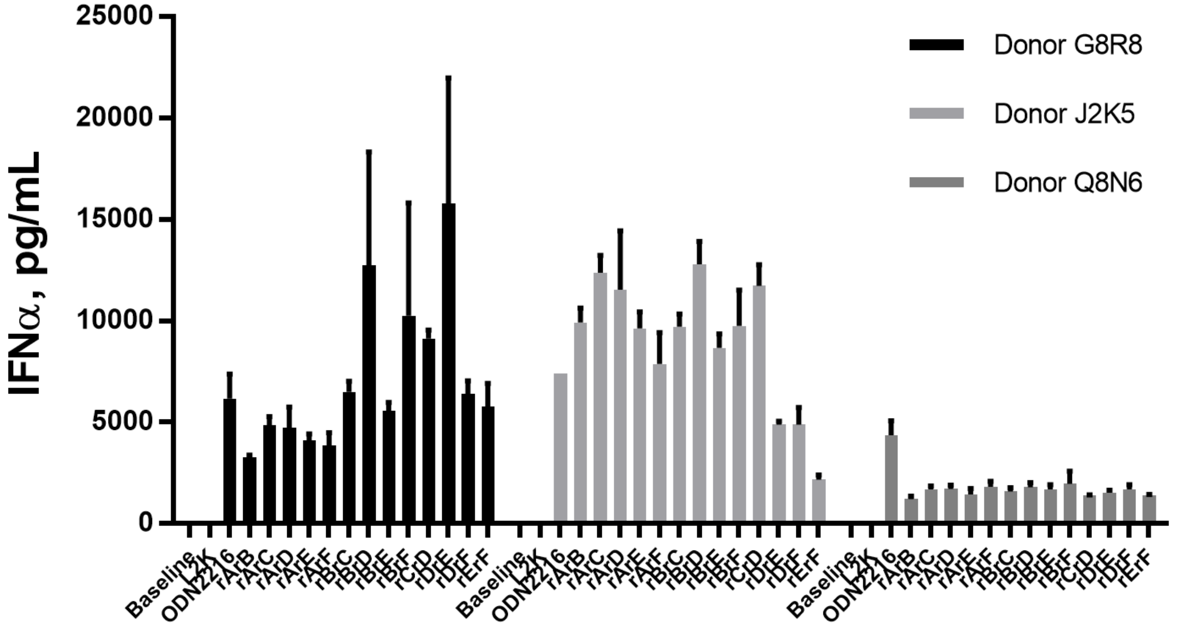


**Figure S1.** IFNα production assessed for PBMC collected from three different donors in response to six –stranded RNA/DNA cubes. The names of the cubes represent the RNA strands present in their compositions. For example, rArB, corresponds to the cube rArBdCdDdEdF.

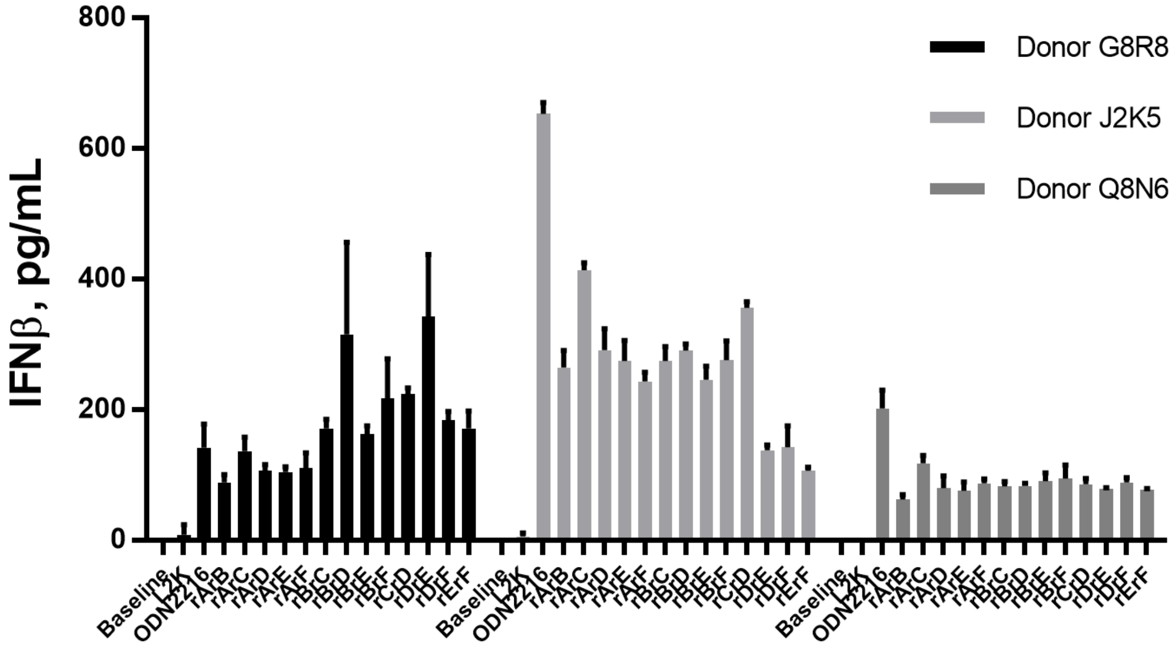


**Figure S2.** IFNβ production assessed for PBMC collected from three different donors in response to six –stranded RNA/DNA cubes. The names of the cubes represent the RNA strands present in their compositions. For example, rArB, corresponds to the cube rArBdCdDdEdF.

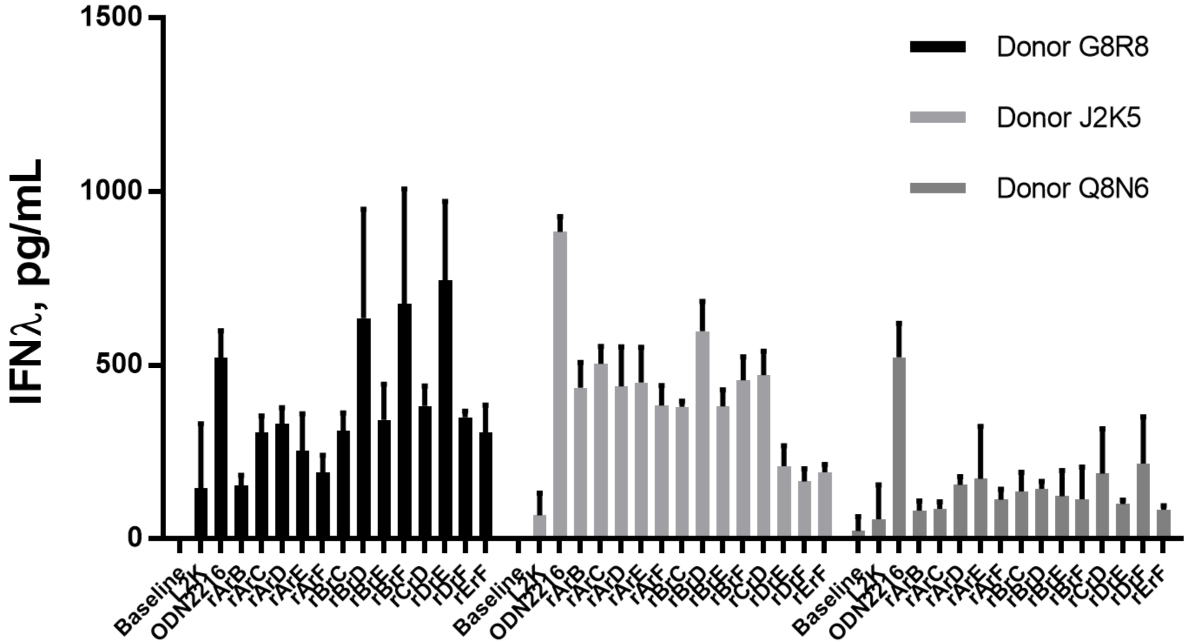


**Figure S3.** IFNλ production assessed for PBMC collected from three different donors in response to six –stranded RNA/DNA cubes. The names of the cubes represent the RNA strands present in their compositions. For example, rArB, corresponds to the cube rArBdCdDdEdF.

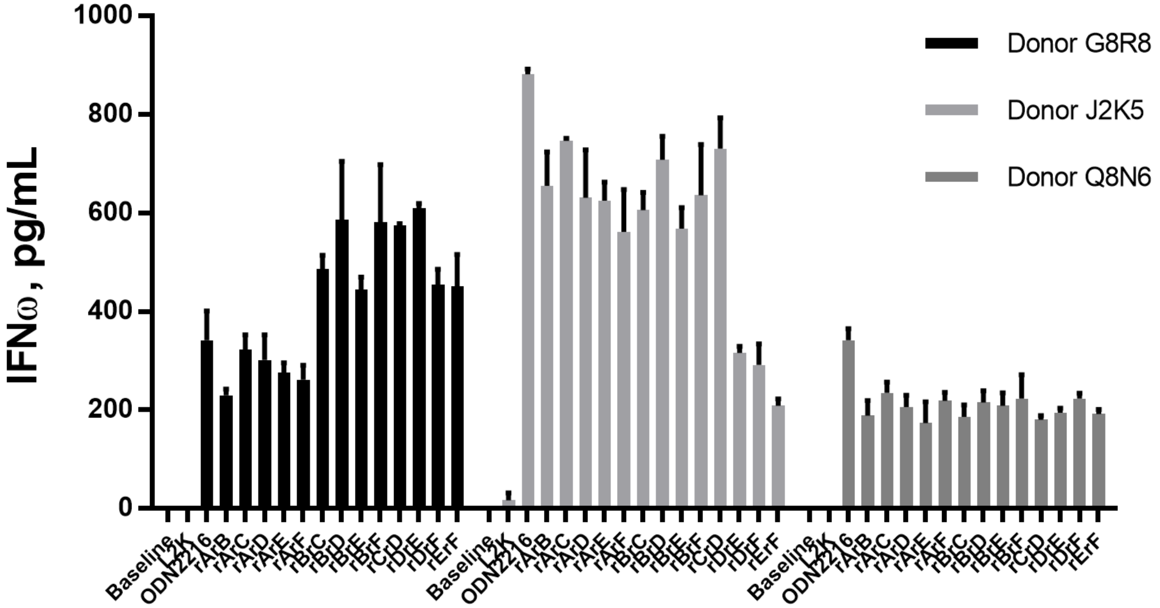


**Figure S4.** IFNω production assessed for PBMC collected from three different donors in response to six –stranded RNA/DNA cubes. The names of the cubes represent the RNA strands present in their compositions. For example, rArB, corresponds to the cube rArBdCdDdEdF.
